# scraps: an end-to-end pipeline for measuring alternative polyadenylation at high resolution using single-cell RNA-seq

**DOI:** 10.1101/2022.08.22.504859

**Authors:** Rui Fu, Kent A. Riemondy, Ryan M. Sheridan, Jay R. Hesselberth, Craig T. Jordan, Austin E. Gillen

## Abstract

Alternative cleavage and polyadenylation (APA) contributes to the diversity of mRNA 3′ ends, affecting post-transcriptional regulation by including or excluding *cis*-regulatory elements in mRNAs, altering their stability and translational efficiency. While APA analysis has been applied broadly in mixed populations of cells, the heterogeneity of APA among single cells has only recently begun to be explored. We developed an approach we termed scraps (<u>S</u>ingle <u>C</u>ell <u>R</u>N<u>A</u> <u>P</u>olyA <u>S</u>ite Discovery), implemented as a user-friendly, scalable, and reproducible end-to-end workflow, to identify polyadenylation sites at near-nucleotide resolution in single cells using 10X Genomics and other TVN-primed single-cell RNA-seq (scRNA-seq) libraries. Our approach, which performs best with long (>100bp) read 1 sequencing and paired alignment to the genome, is both unbiased relative to existing methods that utilize only read 2 and recovers more sites at higher resolution, despite the reduction in read quality observed on most modern DNA sequencers following homopolymer stretches. For libraries sequenced without long read 1, we implement a fallback approach using read 2-only alignments that performs similarly to our optimal approach, but recovers far fewer polyadenylation sites per experiment. scraps also enables assessment of internal priming capture events, which we demonstrate occur commonly but at higher frequency during apoptotic 3′ RNA decay. We also provide an R package, scrapR, that integrates the results of the scaps pipeline with the popular Seruat single-cell analysis package. Refinement and expanded application of these approaches will further clarify the role of APA in single cells, as well as the effects of internal priming on expression measurements in scRNA-seq libraries.

## Introduction

Single-cell sequencing has proven to be an immensely powerful tool for the investigation of cellular heterogeneity in gene expression (scRNA-seq), transcriptional regulation (scATAC-seq), and spatial patterns of gene expression (Visium). However, to date, regulation of gene expression by changes in RNA processing remains understudied, particularly at single-cell resolution. One of the most common alterations to mRNA processing is alternative cleavage and polyadenylation (APA). By altering the length of the 3′ untranslated region (3′ UTR), alternative polyadenylation can remove or introduce regulatory elements recognized by microRNAs (miRNAs) or RNA binding proteins (RBP). Shortly after the advent of widespread high-throughput single-cell sequencing, several groups reported the application of existing informatic approaches designed for extracting APA events from bulk RNA-seq to single-cell libraries (1–3). However, none of these methods approach single-nucleotide resolution, and only one (1) considered the impact of internal priming – priming reverse transcriptase on genomically-encoded A-rich stretches rather than bona fide poly(A) tails – which is widespread in TVN-primed single-cell libraries. We observed that TVN-primed (poly(T) primer with VN anchor at the 3′ end) single-cell RNA-seq libraries, such as the popular 10X Genomics 3′ platform, are fundamentally identical in construction to PAS-seq (4) libraries that measure poly(A) sites at single-nucleotide resolution (Fig. 1A). The “VN” sequence (V is A, G, or C, and N is any base) anchor positions the priming and reverse transcription events at the very start of the poly(A) tail of transcripts, while the number of poly(T)s included in the final library reflects the stretch of Ts in the primer. More recently, several single-cell specific techniques have been reported (5–8), including some that take partial advantage of the single-nucleotide resolution of TVN-primed libraries. However, all of these approaches suffer from one or more of the following issues that reduce their utility: (a) they rely exclusively on gene models for reference, eliminating their ability to identify *de novo* polyadenylation events, (b) they utilize unanchored read 2 alignments (i.e. all read 2 alignments with no requirement for a soft-clipped poly(A) stretch at the 3′ end of the alignment), reducing resolution, (c) they use anchored read 2 alignment, which are relatively rare in these libraries, substantially reducing sensitivity, or (d) they lack a user-friendly, scalable, and reproducible interface integrating with downstream scRNA-seq analysis tools, hampering their adoption. Here, we describe an approach that extracts poly(A) sites from single-cell RNA-seq libraries at near-nucleotide resolution using paired alignments or read 2-only alignments, quantifies them either *de novo* or on an existing reference, and provides an integrated analysis suite for studying APA in single cells.

**Figure 1.**
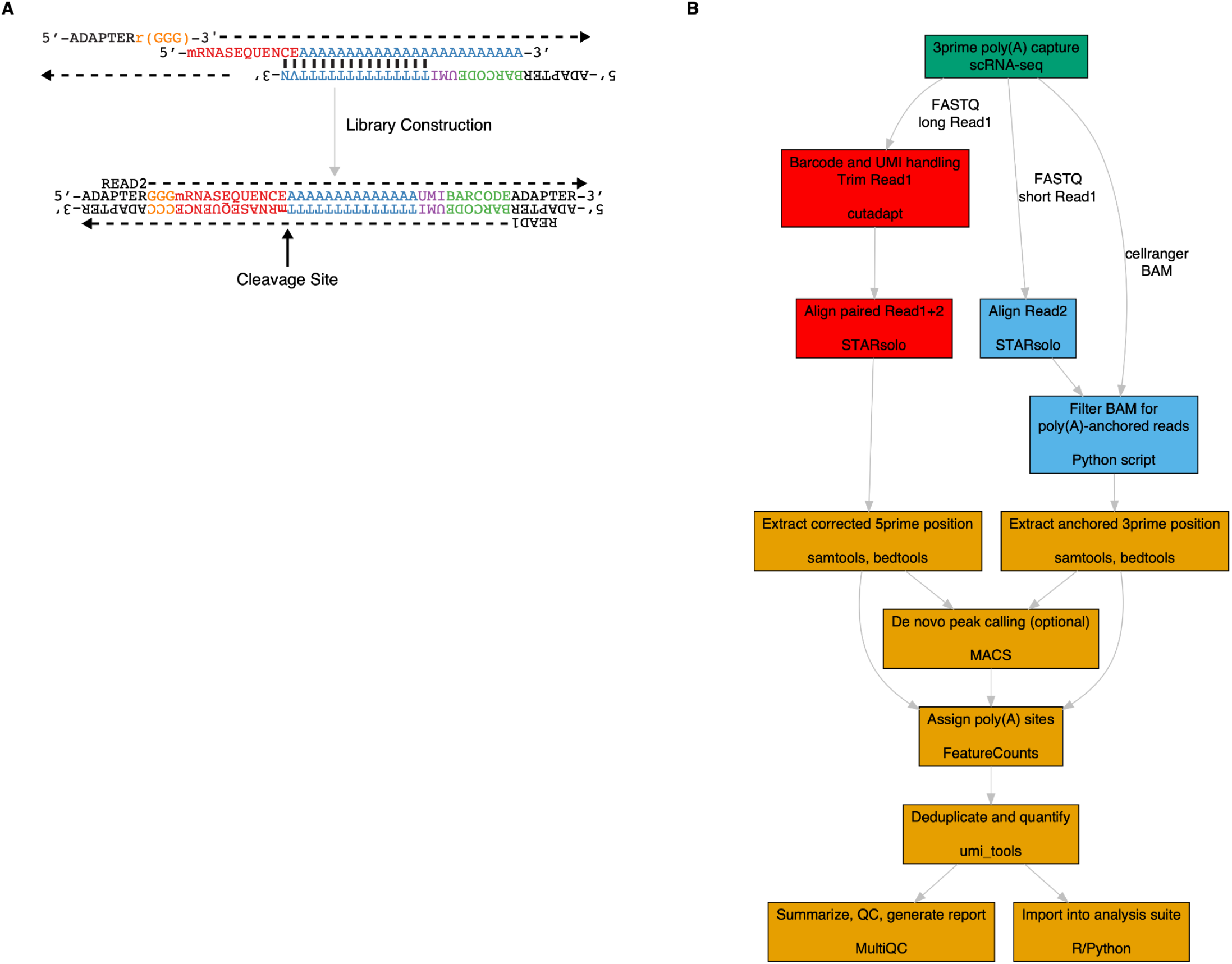
scraps utilizes positional information in TVN-primed libraries to map cleavage and polyadenylation sites at single-base resolution. **A**. Generic library construction schematic for TVN-based assays (PAS-seq, 10X Genomics 3′ single-cell, Drop-seq, etc.). **B.** Schematic representation of the scraps workflow, highlighting the ability to use existing Cell Ranger BAMs, paired alignments (preferred), or new read 2-only alignments to both quantify reference sites and identify novel poly(A) sites.

## Methods

### Software Implementation

For reproducibility and scalability, scraps is implemented as a snakemake (9) pipeline (schematic in Fig. 1B) with conda-powered package management (10). scraps utilizes cutadapt (11) for adapter trimming, STARsolo (12) for genome alignment and UMI/cell barcode handling, featureCounts from subread (13) for feature quantification, umi_tools (14) for UMI deduplication, MultiQC (15) for quality control reporting, and MACS (version 2) (16) for peak calling. Additional functions, including anchored read 2 alignment filtering, are implemented in Python. scrapR is implemented as an R package that integrates with the Seurat single-cell analysis package. scrapR implements the PSI (ψ) calculation described in the LABRAT Python package (17). A full list of dependencies and system requirements, as well as full implementation details, are available in the scraps and scrapR GitHub repositories.

### Paired Read Alignment and Quantification

scraps achieves single-nucleotide resolution mapping of poly(A) sites in single-cell data by using the positional information encoded in read 1 of common TVN-primed libraries (Fig. 1A). This is accomplished by performing a paired-end alignment, rather than the standard read 2-only alignment (in which the paired read 1 is used only for cell barcode and UMI data). This approach requires that read 1 be sequenced to at least 100bp (10X Genomic 3′ libraries; others will vary with regard to minimum length). As many single-cell libraries are sequenced with short read 1 (approximately 94% of all single-cell libraries deposited at SRA), scraps can also use read 2-only alignments, filtered for alignments that end with a soft-clipped poly(A) sequence (i.e. anchored read 2) as other groups have reported (7). However, this fallback method substantially reduces the sensitivity of poly(A) site recovery, as <15% of read 2 alignments contain a soft-clipped terminal poly(A), compared to the 100% of read 1 alignments that contain cleavage site information. While read 1 contains substantially more poly(A) site information than read 2, many short-read DNA sequencers experience significant reductions in read quality following homopolymer stretches independent of overall lane complexity, including the poly(T) stretch in read 1 from 10X Genomics 3′ gene expression libraries. In 62 representative 10X Genomics libraries, we found that while paired alignment produces a predictably lower unique alignment rate than read 2 alone (mean ± sem, paired: 32.9±1.3%, read 2: 64.7±2.4%), substantially more usable alignments are recovered from paired alignments because of the need to filter read 2 for anchored alignments (mean ± sem, paired: 88.0M±13.6M, read 2: 41.0M±8.1M). To confirm that the lower alignment rate observed in paired alignments are not biased relative to the previously reported use of anchored read 2, we compared UMIs recovered at polyA_DB v3 (18) reference sites in all 62 10X Genomics libraries (representative library shown in Fig. 2). UMIs recovered from paired alignments showed high positive correlation with read 2-only (R2 = 0.89, Fig. 2A). Additionally, no significant bias (per-position chi squared p-values all > 0.05) was observed when comparing the 100 bp upstream of all sites in polyA_DB (Fig. 2B) with all sites detectable in paired alignments (Fig. 2C) or read 2-only alignments (Fig. 2D). Similarly, when weighting sequences by the number of UMIs recovered at each site, no significant difference was observed between paired alignments (Fig. 2E) or read 2-only alignments (Fig. 2F), despite a bias toward a relatively small number of highly detectable sequences. Finally, when comparing the overall nucleotide distribution in the 100bp upstream of all sites in polyA_DB with the UMI-weighted sites from paired alignments and read 2-only alignments, no significant bias was observed (Fig. 2G; chi squared p-value > 0.05).

**Figure 2.**
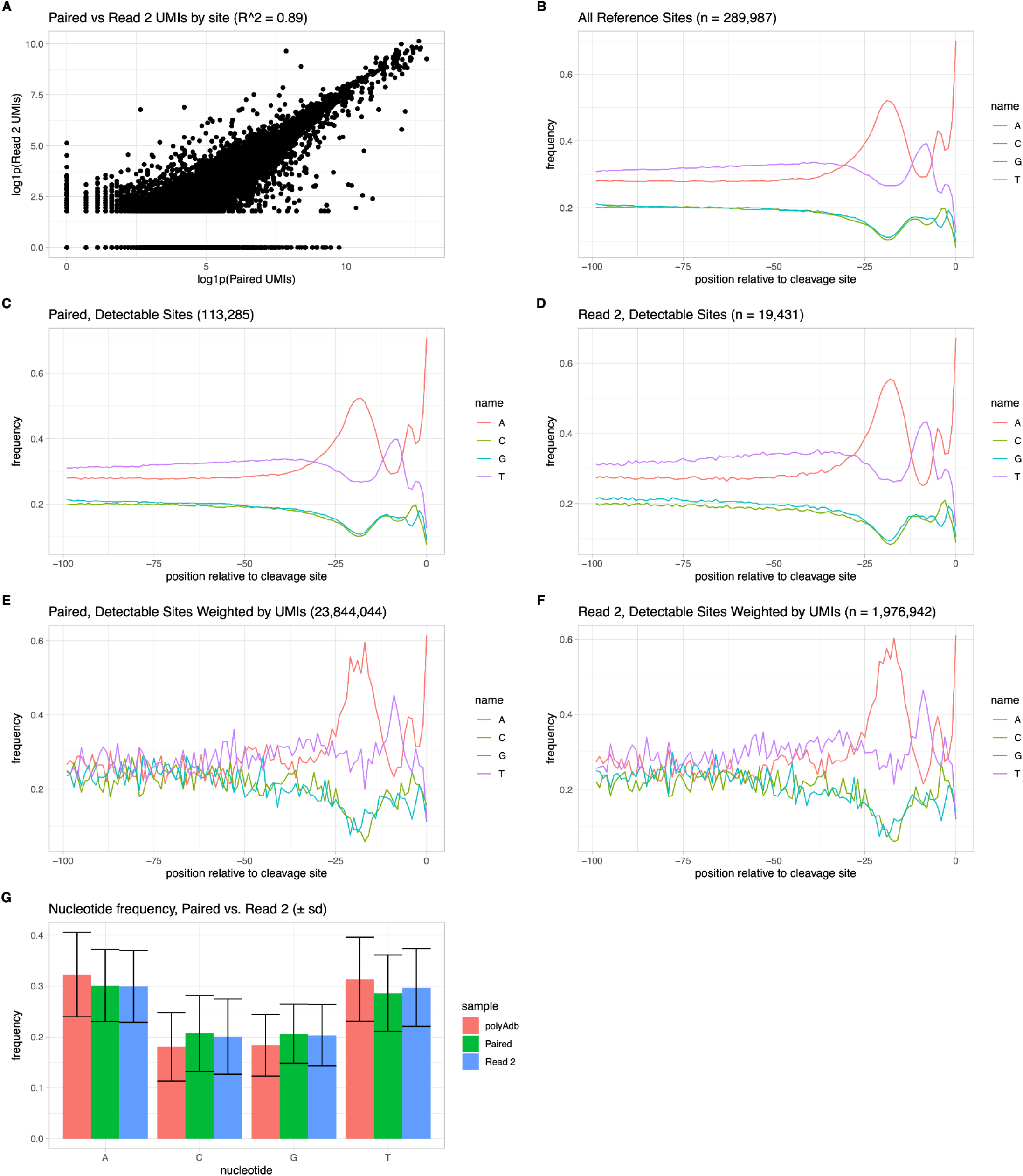
Paired-end alignments with scraps are not sequence biased, despite relatively poor read 1 sequencing performance. **A**. Correlation of UMIs detected using paired (x-axis) and read 2-only (y-axis) alignments for all detectable sites. Pearson R^2^ = 0.89. **B.** Nucleotide frequencies at each position in the 100 basepairs upstream of all poly(A) sites in polyA_DB v3. **C-D.** Nucleotide frequencies in the 100 basepairs upstream of all poly(A) sites from polyA_DB v3 detectable in (**C**) paired and (**D**) read 2-only alignments. **E-F.** Same as C-D, but scaled by the number of UMIs recovered at each site. **G.** Mean nucleotide frequencies in the 100 basepairs upstream of all poly(A) sites in polyA_DB v3 (salmon), sites detectable in paired alignments, scaled by UMIs (green) and sites detectable in read 2-only alignments, scaled by UMIs (blue). p > 0.05 for all comparisons (C vs. D, E vs. F, G).

## Use Cases

### Visualization of APA

scrapR provides three different ways to represent and visualize APA data in single cells. To demonstrate these visualizations, a representative single-cell RNA-seq dataset is shown in Fig. 3. This dataset consists of peripheral blood mononuclear cells from 13 acute myeloid leukemia (AML) patients. As an example, we present an APA event in the NR1D1 (Nuclear Receptor Subfamily 1 Group D Member 1) gene that differentiates AML blasts with a primitive (hematopoietic stem cell-like) phenotype from blasts with a monocytic phenotype. Notably, expression of NR1D1 is known to play a role in the regulation of leukemia stem cells in AML (17). In addition to this biological relevance, we have chosen NR1D1, which is detectable in only 4% of cells in this dataset (Fig. 3C), to highlight that a gene does not need to be highly detectable in order to observe high-confidence APA events. This dataset was generated using the scraps paired-end alignment strategy and quantified using reference sites in polyA_DB v3. As shown in Fig. 3A, two sites - referred from here on as the proximal and distal sites - are detectable in the NR1D1 3′ UTR when plotting scraps alignments in bulk. The dataset is also plotted on a UMAP projection, with cell type clustering and gene-level expression of NR1D1 for reference (Fig. 3B-C). Through the scrapR scraps_to_seurat() function, poly(A) site counts are easily imported and added to Seurat objects as an additional assay. Using the Seurat’s FeaturePlot() function, scraps’ per-cell counts for the proximal and distal sites can be plotted on a UMAP (Fig. 3D-E), showing that the monocytic blasts are heavily biased toward utilization of the proximal site, while primitive blasts are biased towards the distal site. scrapR also implements two ways to represent gene-centric biases in poly(A)-site utilization. The first, referred to as a PSI (ψ) value, is a metric that represents the proximal or distal bias of a gene’s poly(A) site utilization in a single value, independent of the number of poly(A) sites observed in a gene (17). PSI values range from 0 (100% utilization of the most proximal site) to 1 (100% utilization of the most distal site). Fig. 3B shows the PSI values for NR1D1 plotted on the UMAP projection, again highlighting the proximal and distal biases seen in monocytic and primitive blasts, respectively. The second gene-centric metric used by scraps is calculated by fitting a Dirichlet multinomial distribution using the MGLM R package (19) to the per-cell normalized observed counts at each site dataset-wide on a per-gene basis and calculating residuals for each site in each cell. A Dirichlet multinomial model, which accounts for overdispersion, was chosen as it has previously been demonstrated to accurately model count-based mRNA isoform abundance (20). These residuals can be used to identify sites with substantial deviation from the dataset-wide modeled background distribution on a per-cell basis. In Fig. 4G-H, the residuals for the NR1D1 proximal and distal sites are plotted, again highlighting the cell type-specific bias in isoform expression.

**Figure 3.**
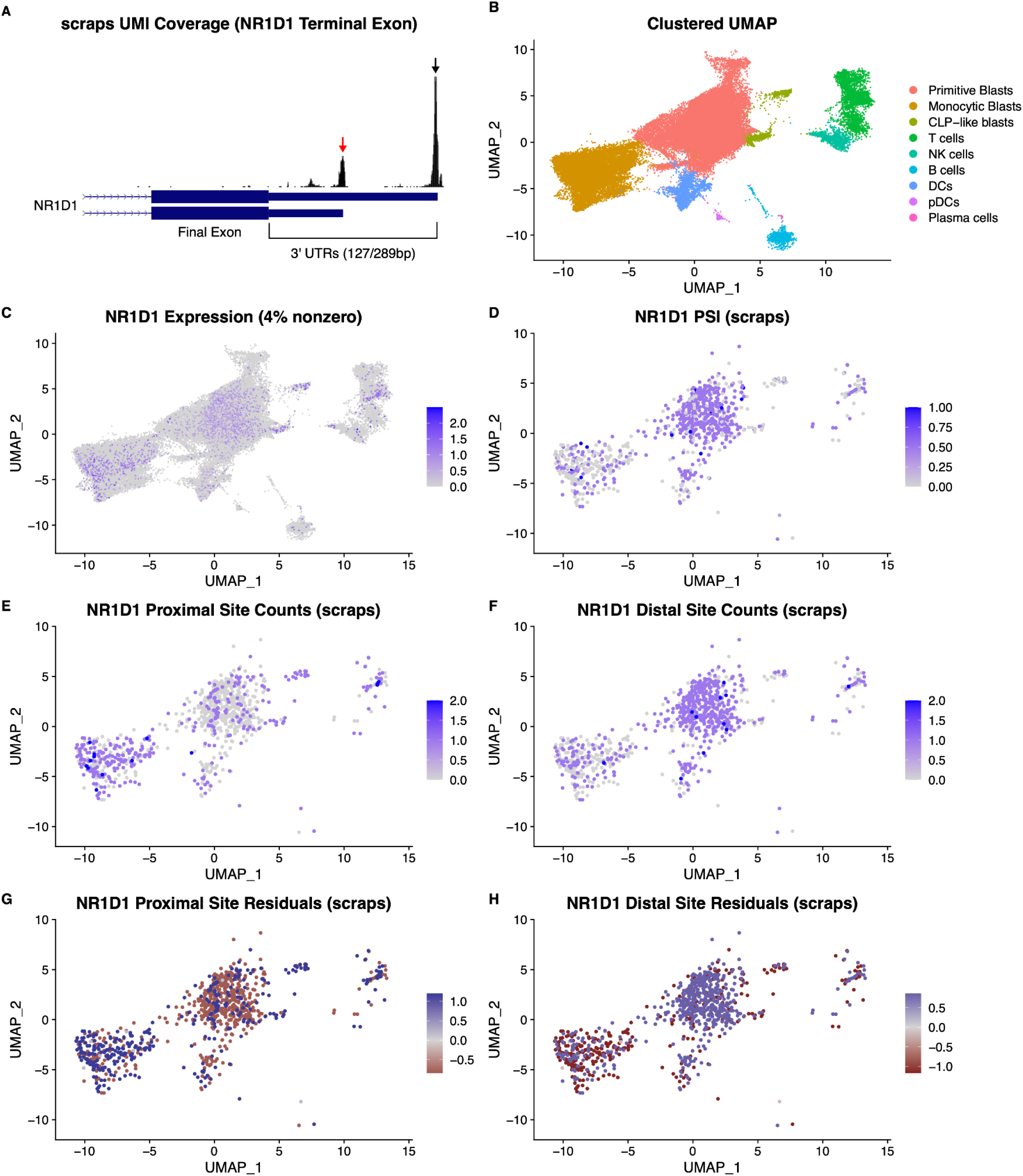
An APA event at NR1D1 discriminates between primitive and monocytic AML blasts. **A**. Bulk UMI coverage demonstrates that two poly(A) sites are detectable in the NR1D1 3′ UTR: the proximal site is indicated with a red arrow, and the distal with a black arrow. A third (unlabeled) peak 5′ of the other two is the result of internal priming. **B-H.** UMAP projections colored by (**B**) cell type, (**C**) NR1D1 RNA expression, (**D**) NR1D1 PSI value, (**E**) NR1D1 proximal poly(A) site counts, (**F**) NR1D1 distal poly(A) site counts, (**G**) NR1D1 proximal poly(A) site Dirichlet multinomial residuals, and (**H**) NR1D1 distal poly(A) site Dirichlet multinomial residuals.

**Figure 4.**
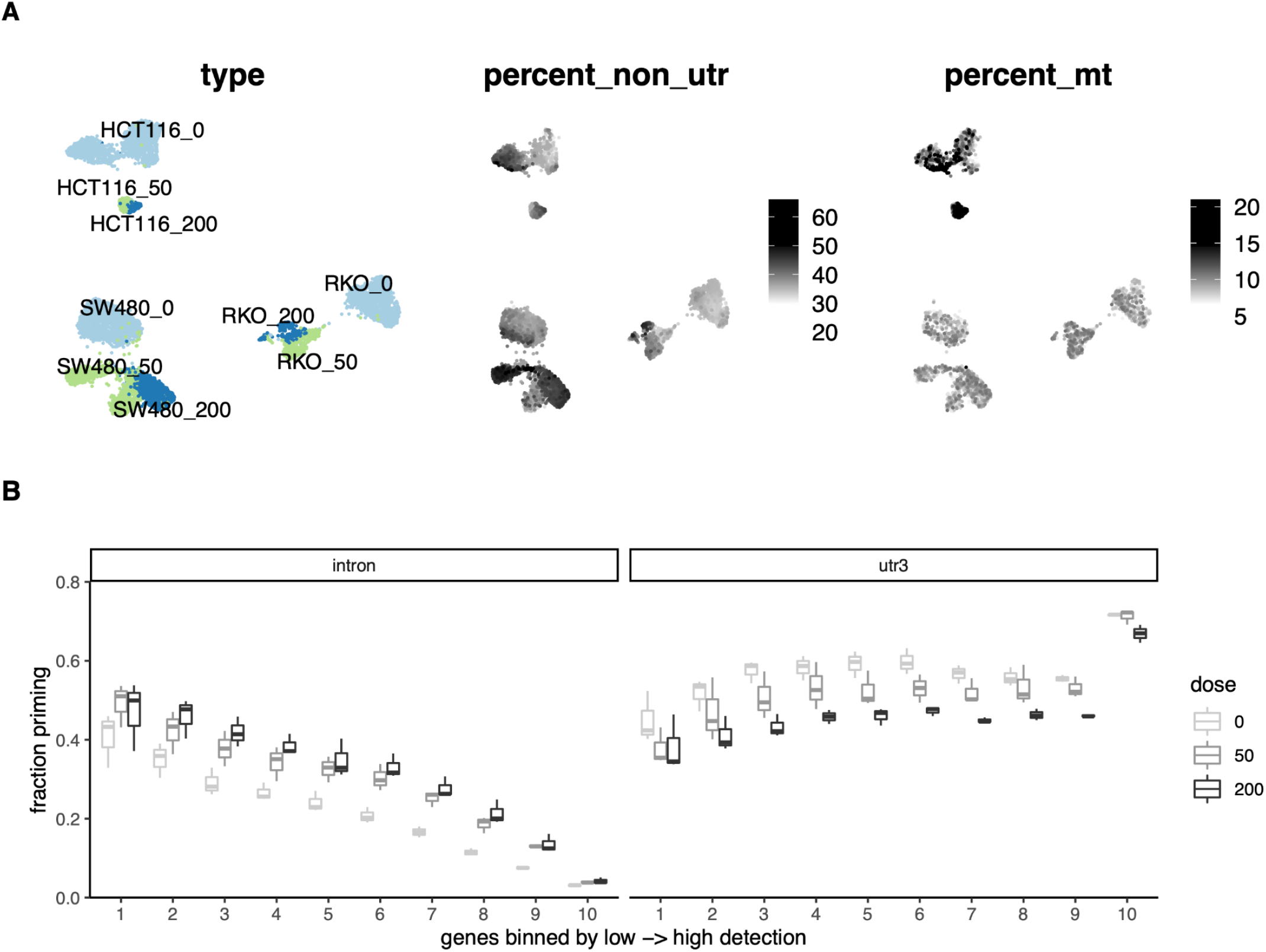
scraps detects increased priming at non-3′ UTR sites in classical apoptosis. **A**. UMAP display of three cell lines and different 5-FU treatment doses, colored by condition (left), overall percentage of UMIs at non-3′ UTR sites determined by scraps(middle), and percentage of cellranger-quantified UMIs that align to mitochondrial genome (right). **B.** Consistent decrease in fraction of priming in the 3′ UTR with increased 5-FU dosage, and corresponding increase in internal (mostly intronic) priming. From scraps quantification, genes were binned by overall UMIs to 10 bins, fraction priming in 3′ UTR or intron was calculated for each bin and cell line, then summarized as boxplots.

### Dimensionality and clustering based on 3′ end isoform expression or APA

While scraps does not itself provide functions to generate new projections or clustering based on the raw site counts, per-gene PSI values, or per-site residuals, each of these matrices can be readily used as input to dimensionality reduction and clustering algorithms. These matrices can also be readily used in place of gene-level count data in multimodal interrogation methods such as Seurat WNN (21), TotalVI (22), and MultiVI (23). In samples with sufficiently deep sequencing and mRNA 3′ end isoform variation, the use of per-site counts allows for isoform-specific projections that may substantially increase resolution over gene expression alone, despite the potential reduction in total UMIs recovered. Similarly, the use of PSI or Dirchlet multinomial residuals allows for the efficient clustering of cells with coordinated changes in mRNA 3′ end isoform utilization. While this is useful for identifying cells with related APA phenotypes, it discards information about relative gene expression and so is not recommended for general projection and clustering without accompanying RNA expression information.

### Detection and filtering of *de novo* poly(A) sites

Due to the fact that scraps accurately maps reverse transcription priming positions, *de novo* cleavage and polyadenylation sites can be identified through peak calling and filtering. scraps optionally uses MACS2 (16) to call peaks which can then be filtered to flag both known poly(A) sites (using polyA_DB, etc.) and peaks derived from internal priming (using CleanUpDtDeq (24), SQANTI (25), or simple upstream sequence filters) to identify novel bona fide cleavage and polyadenylation sites. These sites can then be quantified using the scraps reference-based counting pipeline and incorporated into downstream analyses. This notably allows scraps to be used in organisms without well documented poly(A) sites or robust mRNA isoform information.

### Detection and interpretation of statistically significant APA events

In addition to facilitating the use of standard Seurat statistical tests (i.e. Wilcoxon rank sum) on the per-site Dirichlet multinomial residuals, we also apply DEXSeq, originally designed for bulk RNA-seq differential exon usage, to scraps quantifications to find differential poly(A) site usage between different conditions or cell types. Specifically, only genes with multiple poly(A) sites detected are retained for analysis, and cells are aggregated into pseudo-bulk profiles for each group, either by sample when replicates exist in the study or by random sampling. At thresholds of adjusted p-value < 0.1 and log2 fold-change > 0.25, 3601 PA sites across 2325 genes were discovered between the primitive and monocytic blasts shown in Fig. 3, including both NR1D1 sites. To gain further insights into the regulatory potential of APA events, we annotate statistical testing results described above with differential putative RBP binding sites for each poly(A) site pair of the same gene, from binding likelihood predictions compiled by the oRNAment database (26). The 3′ UTR made from the distal PA site of NR1D1 contains high confidence binding sites of PUM1, KHDRBS3, RBM23, RBMS2, and RBMS3, missing in the isoform using the proximal poly(A) site. PUM1, in particular, is of high interest to understanding monotype maturation and differentiation, since it functions as an RNA decay inducing factor promoting exit of naive pluripotency (27). PUM1 mRNA itself is not found to be differentially expressed between the primitive and monocytic cells (data not shown), however as its only 3′ UTR binding site on NR1D1 mRNA falls in the APA differential region, APA modifies its binding and regulation outcomes. Notably, the proximal and distal poly(A) sites are only separated by 162 basepairs, meaning they would be merged and thus missed in unanchored read 2-only approaches.

### Detection of increased priming at non-3′ UTR sites in classical apoptosis

Accurate mapping of reverse transcription priming positions from scRNA-seq libraries also enables explorations beyond APA. In Fig. 4, we demonstrate scraps-powered confirmation of increased non-3′ UTR priming during classical programmed cell death. Widespread turnover of poly(A) RNA species as a hallmark of classical apoptosis was previously described through biochemical assays and bulk RNA-seq (28). Mitochondrial factor PNPT1 relocates to the cytoplasm during mitochondrial outer membrane permeabilization, triggering the removal of poly(A) tails and then further decay in the 3′ to 5′ direction. At the gene count level, this phenomenon is not detectable in standard single-cell RNA-seq analysis, as a global decrease in mRNA would be “corrected” during library size normalization. However, with decreased length or complete loss of the poly(A) tails, we would predict mispriming events from other sites to increase as apoptosis progresses. Here, we apply scraps to reanalyze an scRNA-seq Drop-seq dataset of human colon cancer cell lines responding to increasing doses of DNA-damage inducing agent 5-FU (29). As expected, though abundant internal priming occurs in all samples as reported previously (30), the percentage of priming events from sites not located in canonical 3′ UTRs (mostly from intronic priming) increases with 5-FU dose in all three cell lines, RKO, SW480, and HCT116, either summarized to each cell after nearest neighbors smoothing (Fig. 4A) or gene expression bins (Fig. 4B). Notably, the percentage of reads from mitochondrial genes, a common metric thought to reflect cell health in scRNA-seq, is inadequate for monitoring apoptosis. By extending scraps functionality to include detection of classical apoptosis (further documented in the supplemental scrapR package), we again demonstrate the power of scraps at revealing previously overlooked RNA biology.

## Summary

Together, scraps and scrapR provide a user-friendly, scalable, and reproducible end-to-end workflow for the identification, quantification, and analysis of single polyadenylation sites at near-nucleotide resolution in single cells using TVN-primed single-cell RNA-seq (scRNA-seq) libraries, including those generated by the 10X Genomics Chromium 3′, Drop-seq, CEL-seq, inDrop, BD Rhapsody, and Microwell-seq protocols. The novel use of paired alignments to take advantage of positional information encoded in read 1 even when base calling quality is suboptimal allows scraps to capture more sites at higher resolution than existing approaches. This high resolution and sensitivity facilitates the identification of novel cleavage and polyadenylation sites that can be quantified in the absence of an existing reference, and further allows for the quantification of sites too close together to be resolved in many existing approaches. While scraps can also quantify sites in libraries lacking long read 1 sequencing, these advantages suggest that future experiments aimed at detecting APA in single cells should be designed to include long read 1 sequencing to ensure optimal resolution and sensitivity. As no additional manipulation at the library prep level is required, scraps is also fully compatible with TVN 3′ capture extension methods such as RNA Capture-seq.

## Data and software availability

The scraps pipeline and scrapR package includes human and mouse reference data derived from polyA_DB v3. These datasets were obtained from the polyA_DB website (https://exon.apps.wistar.org/PolyA_DB/v3/).

Representative scRNA-seq libraries used in Figures 2–3 are fully described in a manuscript currently under review, and a GEO accession will be added here when that manuscript is published.

Single-cell RNA sequencing data from cells undergoing apoptosis was retrieved from GEO accession GSE149224.

Latest scraps source code is available at http://github.com/rnabioco/scraps

Latest scrapR source code is available at http://github.com/rnabioco/scrapr

**License:** MIT license.

**Grant information**: This work was supported by the RNA Bioscience Initiative (funded by a Transformational Research Award from the University of Colorado School of Medicine), United States Department of Veterans Affairs (IK2BX004952-01A1 to AEG), a Leukemia and Lymphoma Society SCOR grant (CTJ), and NIH grants R35CA242376 (CTJ), R35GM119550 (JH), and T32AI074491 (RS).

